# C-terminus CD28 phosphorylation (Y218) modulates IL-2 secretion and antitumor effect of CAR-T cells

**DOI:** 10.64898/2026.01.28.701378

**Authors:** Elena Martinez-Planes, Maria Cecilia Ramello, Miguel G. Fontela, Mohammad-Reza Shokri, Lancia Darville, John Koomen, Youngchul Kim, Daniel Abate-Daga

**Author notes:** Correspondence should be addressed to, Daniel Abate-Daga, PhD, H Lee Moffitt Cancer Center and Research Center Institute Department of Immunology, 12902 USF Magnolia Drive, Tampa, FL 33612.

## Abstract

CD28 is a co-stimulatory component of several second-generation chimeric antigen receptor (CAR)-T cells, providing signals essential for T cell proliferation, survival, and cytokine secretion. However, the specific contribution of individual CD28 intracellular motifs to CAR-T cell function remains incompletely understood. Here, we identify tyrosine 218 (Y218) in the CD28 cytoplasmic domain as a critical regulatory site, and demonstrate that its phosphorylation is essential for optimal CAR-T cell activity. Using a 218F mutant, we show that loss of Y218 phosphorylation leads to impaired IL-2 production and abrogates antitumor efficacy. Transcriptomic profiling of 218F CAR-T cells revealed increased expression of IL-17A, IL-17F, and related cytokines, suggesting a shift toward a pro-inflammatory Th17-like phenotype that may contribute to dysfunction. Mechanistically, we demonstrate that the interleukin-2-inducible T-cell kinase, ITK, mediates Y218 phosphorylation. To further understand the role of this kinase, we engineered a novel CAR incorporating an ITK-binding motif (PYRP), which enhances ITK recruitment, increases Y218 phosphorylation, and boosts IL-2 secretion, and improves anti-tumor efficacy *in vivo*. Our findings underscore the functional relevance of Y218 phosphorylation in modulating CAR-T cell fate and reveal a strategy to fine-tune CAR signaling through targeted kinase recruitment to enhance therapeutic efficacy.

## INTRODUCTION

Chimeric Antigen Receptor T (CAR-T) cell therapy has emerged as a groundbreaking treatment for hematologic malignancies, offering promising clinical outcomes for patients with leukemia, myeloma, and lymphoma ^1–3^. CAR-T cells are designed to replicate aspects of TCR signaling by incorporating intracellular signaling domains from molecules such as CD3ζ and CD28. The CD3ζ domain initiates activation signals upon antigen engagement, while CD28 provides essential costimulatory signals that enhance T cell expansion, survival, and cytokine production—particularly interleukin-2 (IL-2), a key regulator of T cell proliferation and persistence. Although CAR constructs aim to mimic TCR signaling, the extent to which these engineered receptors recapitulate the complex phosphorylation events of the native TCR-CD28 axis remains incompletely understood.

T cell activation is initiated when the T cell receptor (TCR) engages with an antigen-MHC complex on an antigen-presenting cell (APC). This interaction leads to the phosphorylation of immunoreceptor tyrosine-based activation motifs (ITAMs) within the CD3 complex by the Src-family kinase LCK. Phosphorylated ITAMs serve as docking sites for ZAP-70, a tyrosine kinase that, upon activation, phosphorylates the transmembrane adaptor protein LAT ^4–6^. LAT acts as a scaffold protein complex that facilitates the assembly of a signaling hub, attracting proteins such as SLP76, PLCγ, ADAP, NCK, and ITK ^7,8^. ITK becomes activated through phosphorylation by LCK and subsequently phosphorylates PLCγ, leading to its activation. Activated PLCγ hydrolyzes PIP2 into IP3 and DAG, leading to the activation of NFAT, NF-κB and AP-1 transcription factors ^9,10^. These coordinated signaling events result in T cell proliferation, differentiation, and cytokine production, which are essential for an effective immune response.

CD28 is an essential costimulatory molecule that enhances cytokine production and T cell survival. The intracellular tail of CD28 contains three well-characterized motifs: YMNM, PRRP, and PYAP. YMNM includes tyrosine Y191 which, when phosphorylated, promotes binding of PI3K and leads to generation of PIP3. The PRRP motif is a proline-rich site that associates with ITK, which then signals through PLCγ. PYAP subdomain contains both a proline-rich region and a phosphorylated tyrosine (Y209) that initiates signaling by binding LCK ^11,12^.

Through the use of phosphoproteomics, we previously confirmed that tyrosines Y191 and Y209 in the CD28 domain of the CAR are also phosphorylated upon CAR-T cell activation ^13^. Unexpectedly, we also observed robust phosphorylation of the C-terminal tyrosine Y218, a site that remains poorly characterized. This observation suggested a potentially overlooked regulatory mechanism, highlighting the need to explore its contribution to CAR-T cell activity. Here, we demonstrate that Y218 phosphorylation is essential for IL-2 production and anti-tumor effect in a model of pancreatic cancer. Furthermore, we show that promoting ITK recruitment, increases phosphorylation of Y218 and boosts IL-2 production. These findings provide new insights into how TCR-like signaling events can be leveraged to improve CAR-T cell efficacy against solid tumors.

## RESULTS

### CD28 Y218 is rapidly phosphorylated following CAR activation

The CAR construct used in this study targets prostate stem cell antigen (PSCA), which is expressed in various tumor types, including bladder, prostate, and pancreatic cancers ^23^. This CAR construct incorporates a CD28 costimulatory domain and a CD3ζ signaling domain (**Figure 1A**). CAR-T cell generation was performed as outlined in **Figure 1B**.

**Figure 1.**
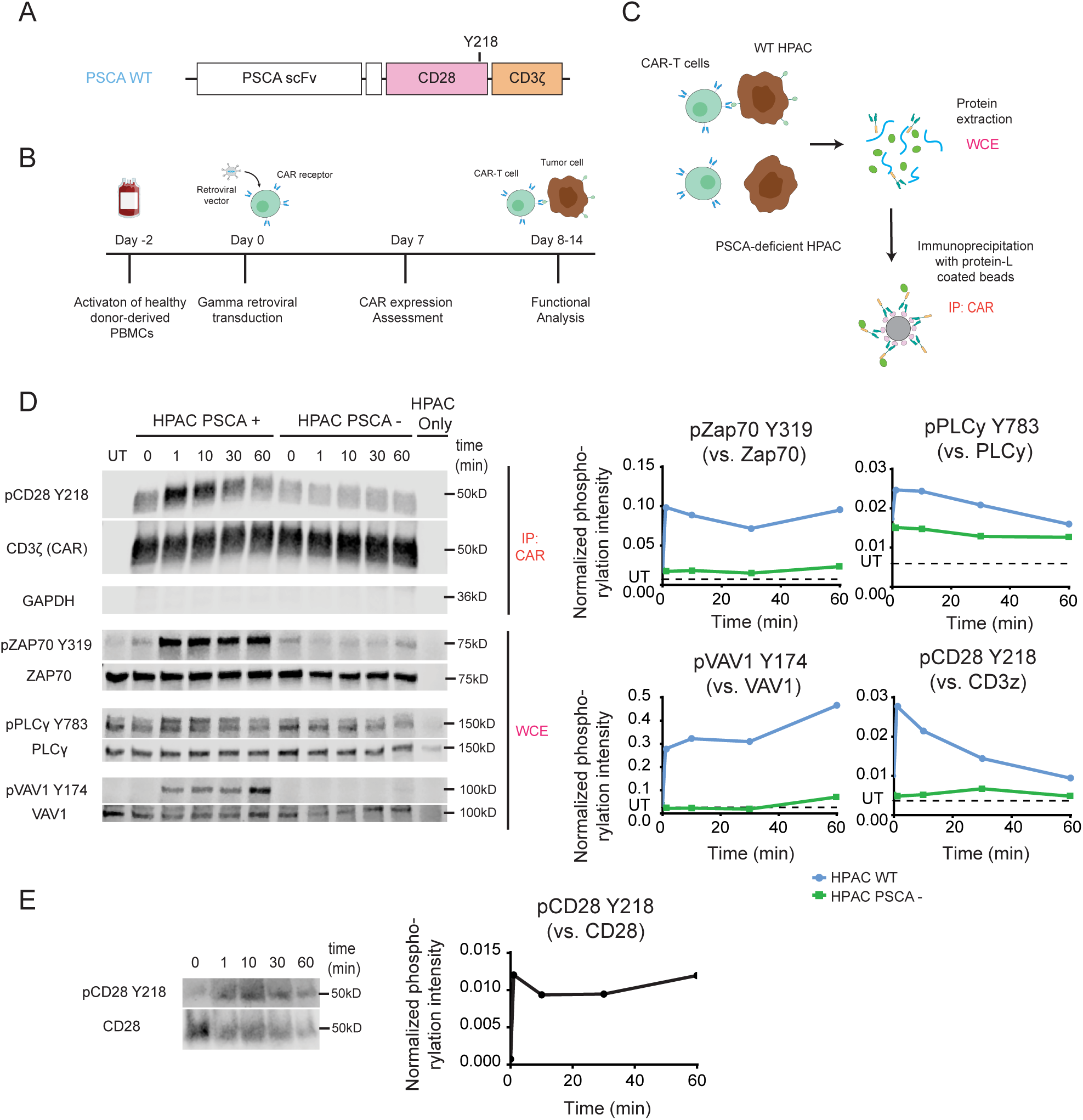
CD28-Y218 is rapidly phosphorylated upon CAR stimulation. **A.** Schematic representation of the CAR construct used in this study: a second-generation CAR receptor containing a CD28 hinge and transmembrane domain, a CD28 costimulatory domain, and CD3ζ signaling domain. **B.** Schematic representation of the protocol used to manufacture CAR-T cells. Healthy donor-derived PBMCs were activated using an anti-CD3 antibody. After 48h, expanded T cells were transduced with a retroviral vector to induce expression of the CAR. Transduction efficiency was assessed 5-7 days after transduction and cells were used for functional analysis between 7-14 days after transduction. **C.** Schematic representation of the protocol used to detect phosphorylation in CAR-T cells. CAR-T cells were cocultured with PSCA-expressing HPAC (HPAC^WT^) or PSCA-deficient HPAC (HPAC^PSCA-KO^) tumor cells for 1, 10, 30 and 60 minutes, followed by protein extraction. For cytoplasmic proteins (ZAP70, PLCγ, VAV1), phosphorylation was detected in the whole cell extract (WCE). For detection of phosphorylated CD28 Y218, the CAR was enriched through immunoprecipitation. **D.** Left Panel. Representative Western blot showing the phosphorylation of CD28 Y218, Zap70 Y319, PLCγ Y783, VAV1 Y174 in PSCA-specific CAR-T cells, following stimulation with HPAC^WT^ or HPAC^PSCA-KO^ cells. Right Panel. Phosphorylation intensity quantified by densitometry (representative plot). CD28 pY218, ZAP70 pY319, PLCγ pY783 and VAV1 pY174 normalized with respect to total CAR (CD3ζ), total ZAP70, total PLCγ and total VAV1 respectively. Untransduced cells (UT) were included as a negative control of immunoprecipitation. **E.** T cells were isolated from PBMCs and stimulated using CD3/CD28 T -activator Dynabeads®. CD28 was then enriched through immunoprecipitation for further analysis. Left Panel. Representative Western blot showing the phosphorylation of endogenous CD28 Y218 at the indicated time points after stimulation. Right Panel. Phosphorylation intensity quantified by densitometry. CD28 pY218 was normalized with respect to total CD28.

In this study, we aim to investigate the role of CD28 Y218 phosphorylation in the context of CAR-T cells. Because phosphorylation events often occur in a tightly regulated sequence during T cell activation, their timing is critical for coordinating downstream signaling. Therefore, we first sought to examine when Y218 phosphorylation occurs relative to other key TCR signaling events. To study phosphorylation kinetics, we stimulated CAR-T cells with wild-type (WT) HPAC cells (HPAC^WT^), which naturally express PSCA. As a negative control, we included PSCA-deficient HPAC cells (HPAC^PSCA-KO^) generated using CRISPR/Cas9 genome editing (**Supplementary Figure 1**). We stimulated cells for 0, 1, 10, 30, and 60 minutes and detected phosphorylation of cytoplasmic proteins (Zap70, PLCγ, and VAV1) directly in the whole cell extract (WCE). To evaluate CD28 phosphorylation, we enriched the CAR by immunoprecipitation (**Figure 1C**).

We found that Y218 phosphorylation occurred rapidly - as early as 1 minute post-stimulation - and gradually declined over the next 60 minutes. This phosphorylation pattern closely resembled that of Zap70 Y319 and PLCγ Y83, which also reached peak intensity at 1 minute post-stimulation. In contrast, VAV1 Y174 phosphorylation remained relatively low during the initial stages and peaked at 60 minutes post-stimulation (**Figure 1D**). Stimulation with HPAC^PSCA-KO^ cells confirmed these phosphorylation events were antigen-specific. These results indicate that Y218 phosphorylation in CAR-T cells represents a very early event in the signaling cascade, comparable to other early phosphorylation events such as Zap70 Y319 and PLCγ Y783.

To determine whether the observed phosphorylation dynamics are specific to the CD28 as a component of the CAR, or reflect a broader regulatory feature of CD28 signaling, we next examined Y218 phosphorylation in the endogenous CD28. To this end, we sorted T cells from PBMCs and activated them with CD3/CD28 T-activator Dynabeads®. We then immunoprecipitated the CD28 molecule and detected phosphorylation using phospho-specific antibodies. We observed that endogenous CD28 also underwent rapid Y218 phosphorylation - within 1 minute post-activation - and maintained similar phosphorylation levels for up to 60 minutes (**Figure 1E**). These results show that Y218 phosphorylation kinetics differ between endogenous CD28 and CAR-integrated CD28. Although phosphorylation occurred rapidly in both contexts, it persisted in endogenous CD28 but declined progressively in the CAR construct. This difference in phosphorylation dynamics may be influenced by variations in ligand-receptor interaction strength between CAR and TCR signaling and/or may indicate distinct functional roles of CD28 Y218 in CAR versus endogenous CD28 signaling.

### Y218F mutant CAR-T cells display lower IL-2 secretion and impaired antitumor activity in a model of pancreatic cancer

To evaluate the functional role of the CD28-Y218 phosphorylation in the context of CAR-T cells, we generated a mutant CAR containing a non-phosphorylatable phenylalanine residue in position 218 (**Figure 2A**). We detected no significant effect of the 218F mutation in CAR expression, nor did we detect changes in the CD4/CD8 ratios (**Supplementary Figure 2A**).

**Figure 2.**
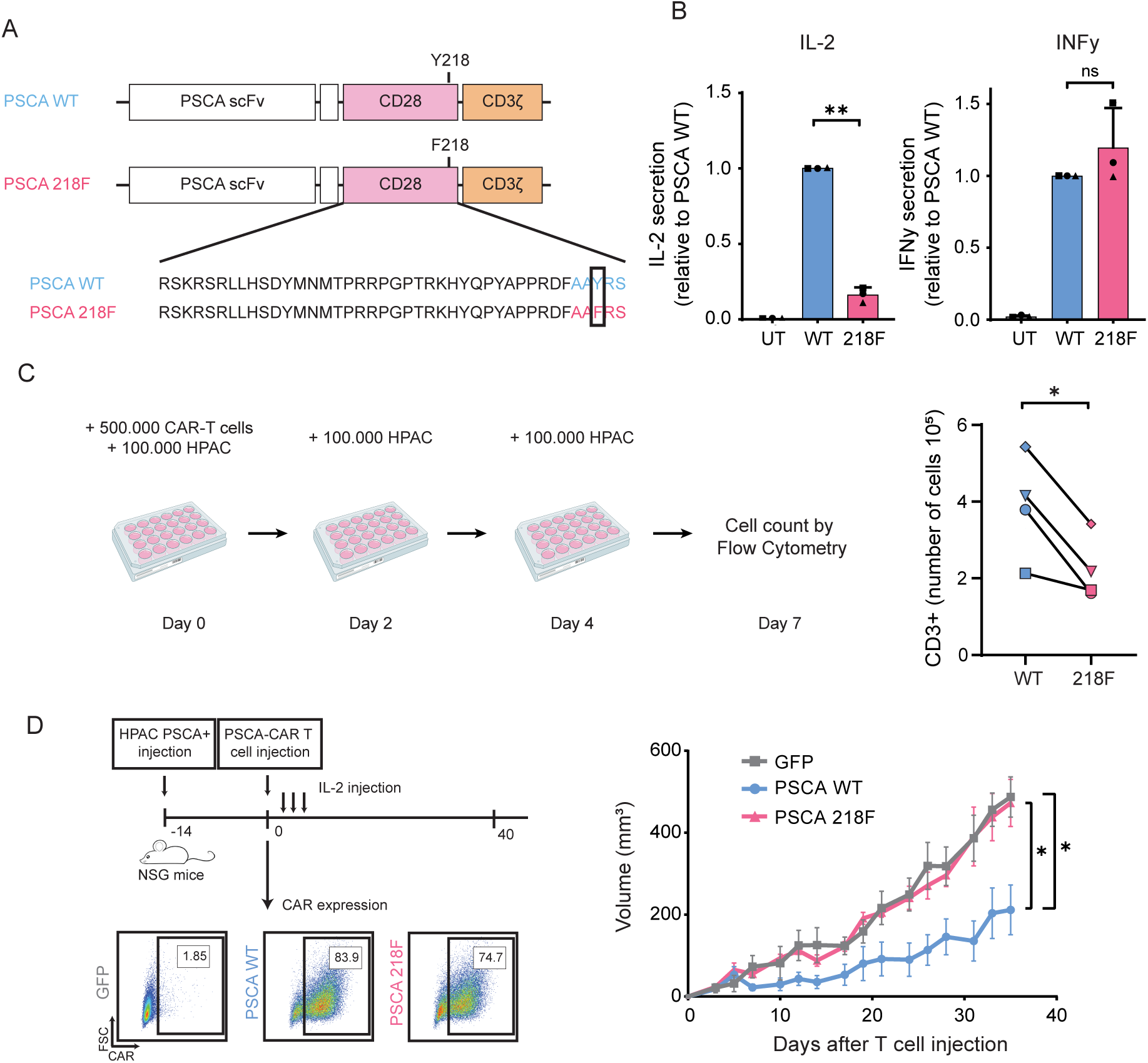
Y218F mutation in PSCA CAR-T cells prevents IL-2 secretion and impairs antitumor activity in a model of pancreatic cancer. **A**. Schematic representation of second-generation chimeric antigen receptors targeting PSCA. The 218F mutant construct contains a point mutation in the AAYRS motif (the tyrosine (Y) is substituted with phenylalanine (F)). **B**. PSCA-targeted WT and 218F CAR-T cells were cocultured with HPAC^WT^ cells at a 1:1 ratio. Supernatant was collected after 24 hours and cytokine production was measured by ELISA. Cytokine levels were normalized to that of WT CAR. Significance was determined by t-test with Welch’s correction. ** = P<0.01. Each symbol represents an independent experiment from 3 different healthy donors. Data is represented as the mean ± standard deviation (SD). **C**. Left panel. Schematic representation of the experimental design. CAR-T cells were cocultured with HPAC^WT^ tumor cells at a 5:1 ratio. On days 2 and 4, 100.000 HPAC^WT^ cells were added to the coculture and on day 7, cells were collected to be quantified. Right Panel. Number of CD3^+^ cells collected on day 7 after repetitive antigen stimulation. Significance was determined by paired t test. * = P<0.05. Each symbol represents an independent experiment from 4 different healthy donors. **D.** Left Panel. Schematic representation of the experimental design. NSG mice were injected subcutaneously with HPAC^WT^ cells and 14 days after tumor injection, PSCA WT or 218F CAR-T cells were injected intravenously and tumor volume was measured for 35 days. Lower Left panel: CAR expression 1 day before adoptive cell transfer. Right panel: Tumor volume (mm^3^) was monitored for 35 days (n=5). Representative example of two independent experiments. Significance was determined by linear regression and one-way ANOVA. * = P<0.05.

To assess the impact of the 218F mutation on their cytolytic capacity, we compared the *in vitro* cytotoxicity of WT and 218F CAR-T cells using the xCELLigence real-time cytotoxicity assay (RTCA). Both WT and 218F CAR-T cells induced rapid and robust killing of HPAC^WT^ target cells within 20 hours (∼80% cytolysis), while untransduced T cells exhibited minimal killing. Quantification of killing at 24 hours post T cells addition revealed substantial donor variability but no significant difference between WT and 218F CAR-T cells. These results indicate that phosphorylation at Y218 is dispensable for *in vitro* cytotoxicity (**Supplementary Figure 2B**). To explore whether the 218F mutation affects cytokine release upon antigen stimulation, we cocultured WT and 218F CAR-T cells with HPAC^WT^ target cells for 24 hours and measured cytokine levels in the supernatant. 218F CAR-T cells produced markedly less IL-2 compared to their WT counterparts, with an average fold change of 0.16, despite donor variability (P=0.0012). In contrast, IFNy secretion remained comparable between WT and 218F CAR-T cells, with no significant difference observed (**Figure 2B**). To assess whether IL-2 production was impaired in CD4⁺ or CD8⁺ T cells, we separated these two T cell subsets using CD4 and CD8 MicroBeads. We then cocultured CAR-T cells with HPAC^WT^ cells at varying CD4:CD8 ratios and measured IL-2 secretion. As expected, CD4⁺ T cells were the primary source of IL-2. The difference in IL-2 production between WT and 218F CAR-T cells was observed exclusively in the CD4⁺ subset, with no detectable difference in CD8⁺ T cells. Consequently, in mixed populations, higher CD4:CD8 ratios amplified the difference in IL-2 production between WT and 218F CAR-T cells, whereas low CD4:CD8 ratios minimized this difference (**Supplementary Figure D**). These results suggest that donor-to-donor variability in CD4:CD8 composition may contribute to the heterogeneity in IL-2 production observed across experiments. Because IL-2 is critical for T cell differentiation, survival, and proliferation ^24^, we next examined whether the reduced IL-2 production observed in 218F CAR-T cells impaired their proliferative capacity. To do so, we stained CAR-T cells with CellTrace™ Violet to track cell division over time. As expected, 218F CAR-T cells proliferated to a lesser extent than their WT counterparts (**Supplementary Figure C**). To evaluate this effect under more physiologically relevant conditions, we examined the ability of WT and 218F CAR-T cells to survive and proliferate under repetitive antigen stimulation, mimicking chronic exposure in the tumor microenvironment. We co-cultured CAR-T cells with HPAC^WT^ cells and added fresh tumor cells every two days to maintain antigen exposure. On day 7 (after three rounds of stimulation), we collected the cells and quantified their number. Consistently, we recovered fewer 218F CAR-T compared to the WT (**Figure 2C**), suggesting that the mutation impairs CAR-T cell expansion and/or persistence over time. We next compared the in vivo performance of WT and 218F CAR-T cells using a pancreatic cancer xenograft model. We injected 5×10^6^ CAR-T via tail vein injection into NSG mice bearing subcutaneous HPAC^WT^ tumors. Tumor volume was monitored for 35 days. WT CAR-T cells significantly delayed tumor progression, with reduced tumor growth evident by day 8 and sustained over the course of the experiment. In contrast, tumors in mice treated with 218F CAR-T progressed at rates similar to those in the mock-transduced group, showing no measurable antitumor activity (**Figure 2D**).

Taken together, our findings identify Y218 phosphorylation within the CD28 costimulatory domain as a critical regulator of CAR-T cell function. Although Y218 phosphorylation was not necessary for IFNγ production or in vitro cytolytic activity, it proved critical for IL-2 secretion, as well as for T cell proliferation and expansion during chronic antigen stimulation. Consistent with these roles, the 218F mutation led to a complete loss of antitumor efficacy *in vivo*.

### Differences between WT and 218F CAR-T cells are found at both signaling and transcriptomic levels

Upon engagement of the CAR receptor, several tyrosine, serine, and threonine residues undergo phosphorylation. These phosphorylated residues serve as docking sites for enzymes and adaptor proteins, facilitating the assembly of signaling complexes that initiate downstream signaling cascades. To elucidate the mechanism underlying the impaired antitumor efficacy of the 218F mutant, we investigated how the 218F mutation affects phosphorylation dynamics and the recruitment of proteins to the CAR receptor following antigen engagement. CAR-T cells were stimulated with tumor cells for 0, 1, 10, 30, and 60 minutes, after which phosphorylation was assessed using phospho-specific antibodies. Phosphorylation of several early proteins (CD3ζ Y142, ZAP70 Y319 and VAV1 Y174) were unaffected by the ablation of Y218 phosphorylation (**Supplementary Figure 3A**). In contrast, we observed a consistent increase in SLP76 S376 phosphorylation following stimulation. Although phosphorylation levels were very similar at early time points, the two curves began to diverge at 10 minutes post-stimulation. The most apparent difference was observed at 60 minutes (P-value = 0.0278) (**Figure 3A**).

**Figure 3.**
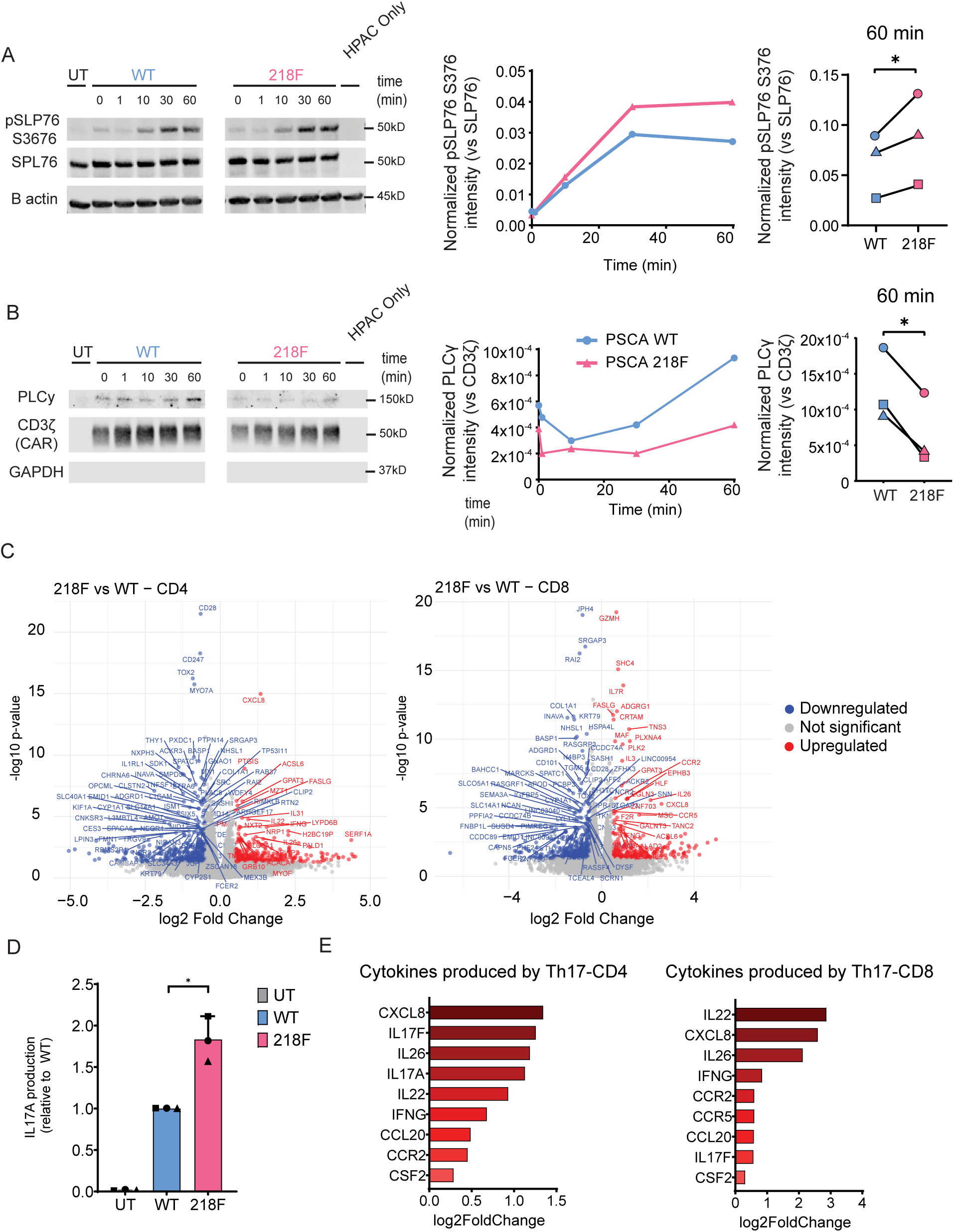
CD28 Y218 phosphorylation regulates PLCγ and SLP76 signaling and drives changes in the Th17 transcriptomic profile. **A-B**. PSCA WT and 218F CAR-T cells were cocultured with HPAC^WT^ cells for 0, 1, 10, 30 or, 60 min. **A.** After protein extraction, CAR-associated proteins were co-immunoprecipitated using protein-L coated beads. PLCγ abundance in the immunoprecipitate was analyzed by Western blot. Left panel: Representative Western blot. Right panel: Normalized CAR-bound PLCγ intensity quantified by densitometry (representative plot), and CAR-bound PLCγ 60 min post-stimulation, showing 3 biological replicates corresponding to different donors, analyzed in independent experiments. Densitometry values were normalized to total CAR (CD3ζ) **B.** SLP76 pY376 was analyzed using phospho-specific antibodies. Left panel: Representative Western blot of SLP76 pY376 in CAR-T cells after stimulation with HPAC^WT^ cells. Right panel: Normalized SLP76 pY376 intensity quantified by densitometry (representative plot) and SLP76 pY376 at 60 min post-stimulation. Each symbol represents an independent experiment from 3 different healthy donors. Untransduced cells (UT) were included as a negative control of immunoprecipitation. Densitometry results were normalized relative to total SLP76. Significance was determined by paired t-test. * = P<0.05. **C and E**. Transcriptomic profile of WT vs 218F CAR-T cells. CAR-T cells were cocultured with HPAC^WT^ cells for 24 hours and subsequently sorted into CD4⁺ and CD8⁺ subsets using the MACSQuant® Tyto® system prior to RNA extraction. **C.** Volcano plot representing the genes that are differentially expressed in 218F CAR-T cells vs WT CAR-T. CD4+ cell is the left, CD8+ cell in the right. **D.** CAR-T cells were cocultured with HPAC^WT^ target cells and supernatant was collected 24hours hours after. IL-17A production was analyzed by ELLA. Cytokine production of 218F and UT was normalized to WT CAR. Significance was determined by t-test with Welch’s correction. * = P<0.05. Each symbol represents an independent experiment from 3 different healthy donors. Data is represented as the mean ± standard deviation (SD). **E.** Gene expression of cytokine-related genes associated with Th17 cells. Bar plot shows log₂ fold change in gene expression of 218F CAR-T cells relative to WT CAR-T cells. CD4⁺ cells are shown on the left and CD8⁺ cells on the right.

In addition, we evaluated how the 218F mutation affects binding of different proteins to the CAR. CAR-T cells were stimulated with target cells for 0, 1, 10, 30 and 60 minutes and then, CAR and CAR-binding proteins were immunoprecipitated using protein-L beads. We observed a time-dependent accumulation of proteins bound to the CAR, peaking at 60 minutes post-stimulation. The binding of most of those proteins (LCK, ZAP70, PLCγ, ITK) was not affected by the 218F mutation (**Supplementary Figure 3B**). However, PLCγ association with the CAR was consistently reduced in the absence of Y218 phosphorylation. Following stimulation, CAR-bound PLCγ increased in WT CAR, reaching a peak at approximately 60 minutes. In contrast, CAR-bound PLCy in the 218F CAR remained relatively unchanged compared to non-stimulation levels. The difference between WT and 218F is most apparent at 60 minutes post-stimulation (P-value = 0.012) (**Figure 3B**). PLCγ is recruited to the signalosome via direct binding with LAT ^7^ and it regulates Calcium and DAG signaling, leading to activation of the transcription factor NFAT and the kinase PKC.These results suggest that the Y218 phosphorylation of CD28, rather than having a general effect on signaling, is involved in a specific complex signaling pathway that involves SLP76 and PLCγ signaling.

To further investigate the mechanisms underlying the differences observed between WT and 218F, we performed transcriptomic profiling via RNA sequencing (RNA-seq). WT and 218F CAR-T cells were stimulated with HPAC^WT^ target cells for 24h. Following stimulation, CD4^+^ and CD8^+^ T cells were isolated using MACSQuant Tyto technology, and total RNA was extracted and sequenced. Principal component analysis (PCA) revealed that the major source of variance was attributable to T cell subset identity - CD4⁺ vs CD8⁺. However, when CD4⁺ and CD8⁺ cells were analyzed independently, clear transcriptional differences emerged between WT and 218F CAR-T cells within each subset (**Supplementary Figure 3C**). Volcano plots summarizing differentially expressed genes (DEGs) for CD4⁺ and CD8⁺ compartments are shown in **Figure 3C**. DEGs were subsequently subjected to pathway enrichment analysis using Ingenuity Pathway Analysis (IPA). IPA highlighted significant alterations in cytokine-related signaling pathways between WT and 218F CAR-T cells, including “Role of Cytokines in Mediating Communication between Immune Cells”, “Interleukin-4 and Interleukin-13 Signaling”, “Interleukin-10 Signaling”, and “IL-27 Signaling” — observed in both CD4⁺ and CD8⁺ populations. Notably, CD4⁺ T cells showed downregulation of genes involved in the Th2 signaling pathway, suggesting a potential reshaping of the CD4⁺ compartment. Interestingly, “IL-17 Signaling” also emerged as a significantly enriched pathway in both CD4⁺ and CD8⁺ cells, with a z-score of 1.732 and p-value of 6.46 × 10⁻_. In CD4⁺ T cells, we observed upregulation of IL17A, IL17F, and IL-17-associated genes (**Supplementary Figure 3D**). To validate these findings, we measured IL-17 secretion in supernatants of CAR-T cells stimulated with tumor cells for 24h. Consistent with the RNA-seq results, 218F CAR-T cells secreted significantly more IL-17A protein than WT, with an average 1.8-fold increase (P = 0.035) (**Figure 3D**). We also identified RAR related Orphan Receptor C (RORC, encoding RORγt) as a key upstream regulator of the changes observed in CD4⁺ cells (activation z-score = 2.14; p = 1.34×10⁻_). RORγt is the lineage-defining transcription factor for Th17 cells, promoting their differentiation and expression of Th17-associated genes ^25,26^.

In the CD8⁺ subset, although IL-17 cytokine expression itself was not significantly upregulated, DEGs enriched in the “IL-17 Signaling” pathway were still observed, suggesting that CD8⁺ cells may respond to increased IL-17 production by CD4⁺ cells (**Supplementary Figure 3B**). Supporting this, both CD4⁺ and CD8⁺ cells showed increased expression of additional Th17-associated cytokines (**Figure 3E**), including IL-26 and IL-22, which are typically co-expressed with IL-17 in Th17 cells. Collectively, these findings indicate that the 218F mutation alters the cytokine production landscape of CAR-T cells, promoting a Th17-skewed phenotype in the CD4⁺ compartment, characterized by upregulation of IL-17 and related effector molecules.

### ITK is required for CD28-Y218 phosphorylation

Our results highlight the critical role of CD28 Y218 phosphorylation in CAR-T cell activity and anti-tumor effects, although the regulatory mechanisms of this site remain unclear. Therefore, we next sought to identify the kinase responsible for this phosphorylation event. Previous *in silico* predictions ^13^ pointed to Interleukin-2-Inducible T-cell Kinase (ITK) as the main candidate. To investigate the role of ITK in CD28 Y218 phosphorylation, we employed two complementary approaches: (1) pharmacological inhibition and (2) genetic ablation of ITK. In the first approach, we inhibited ITK activity using three distinct ITK inhibitors: BMS-509744, Ibrutinib and GNE-9822. CAR-T cells derived from primary T cells were incubated with ITK inhibitors at 1μM, as previously described ^15^, for 2 hours. Following pre-incubation, CAR-T cells were stimulated with HPAC^WT^ for 10 minutes. As expected, stimulation with the target cells induced phosphorylation of CD28 Y218, but this phosphorylation was consistently reduced in cells pre-treated with any of the three ITK inhibitors (**Figure 4A**). In the second approach, we established a clonal population of ITK-deficient Jurkat cells (**Supplemental Figure 4A**). The CAR construct was then transduced into both WT and ITK-KO Jurkat cells, achieving comparable transduction efficiencies as observed in primary T cells (**Supplemental Figure 4B**). CAR-expressing Jurkat cells were activated with HPAC^WT^ cells for 10 minutes. Stimulation of CAR-expressing WT Jurkat cells resulted in robust CD28 Y218 phosphorylation, similar to the pattern observed in primary CAR-T cells. In contrast, CD28 Y218 phosphorylation was absent following stimulation of CAR-expressing ITK-KO Jurkat cells (**Figure 4B**). Together, these results demonstrate that both small-molecule inhibition and genetic deletion of ITK impair CD28 Y218 phosphorylation, strongly implicating ITK as a key mediator of this phosphorylation event.

**Figure 4.**
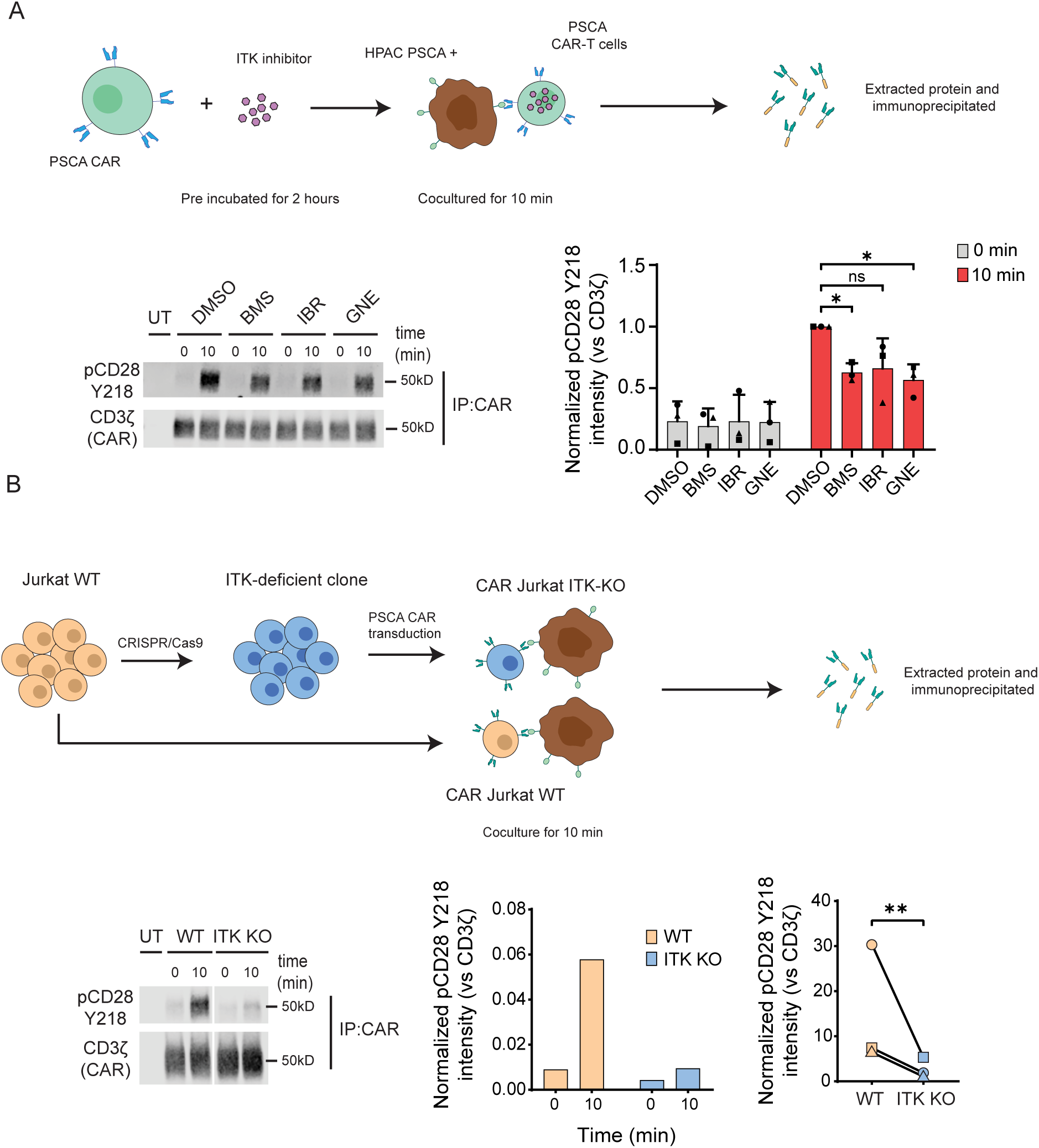
ITK is required for CD28 phosphorylation. **A**. Top panel: Schematic representation of the protocol used to inhibit ITK activity. PSCA CAR-T cells were pre-treated for 2 hours with ITK inhibitors 1μM (BMS-509744, Ibrutinib or GNE-9822) or DMSO as control and then cocultured with HPAC^WT^ cells for 0 or 10 minutes. Left panel: Representative Western blot of CD28 pY218 in CAR-T cells after stimulation with HPAC^WT^ cells. Right panel: Normalized CD28 pY218 intensity with respect to CAR (CD3ζ). Significance was determined using one-way ANOVA. * = P<0.05. Each symbol represents an independent experiment from 3 different healthy donors. Data is represented as the mean ± standard deviation (SD). **B.** Top panel. Schematic representation of the experimental protocol. ITK expression was disrupted using CRISPR/Cas in Jurkat cells. WT and ITK-KO Jurkat cells were transduced to express a PSCA-specific CAR and cocultured with HPAC^WT^ cells for 0 or 10 min. Left panel. Representative membrane of CD28 pY218 in CAR-T Jurkat cells after stimulation. Right panel. Normalized CD28 pY218 intensity quantified by densitometry (representative plot) and CD28 pY218 at 10 min post-stimulation. Significance was determined by paired t-test. ** = P<0.01. Each symbol represents one of 3 independent experiments, performed with different donor T cells.

### PYRP mutation increases IL-2 secretion and improves antitumor efficacy

Given the dependency of CD28 Y218 phosphorylation on ITK, we hypothesized that redirecting ITK to the CAR could promote CD28 Y218 phosphorylation. To enhance ITK binding to the CAR, we generated a novel mutant CAR construct (PYRP) containing an additional synthetic ITK-binding motif within the CD28 domain (**Figure 5A**). ITK contains a Src Homology 3 (SH3) domain that binds consensus PXXP motifs within proline-rich regions ^27^ and it has been reported to bind a variety of proteins including Sam68, WASP ^28^, VAV1, TSAD ^29^ and SLP76 ^30^. In this mutant, alanine (A) 217 and serine (S) 220 were substituted with proline (P), to generate a proline-rich domain that mimics the natural ITK-binding site. We then tested ITK binding to the mutant PYRP CAR by stimulating WT and PYRP CAR-T cells with HPAC^WT^ cells for 0, 1, 10, 30, and 60 minutes. Following stimulation, we immunoprecipitated the CAR using Protein L–conjugated beads and analyzed the presence of ITK in the co-immunoprecipitate. In both WT and PYRP CAR-T cells, CAR-bound ITK was low at early time points but increased substantially at 60 minutes. Differences between WT and PYRP were more apparent at this later time point, with more ITK detected in complex with the PYRP CAR compared to the WT **Figure 5B**. We then evaluated the effects of the PYRP mutation in CD28 Y218 phosphorylation. WT and PYRP CAR-T cells were cocultured with HPAC^WT^ cells for 10 minutes, and the CAR was immunoprecipitated. Because the PYRP mutation alters the conformation of the epitope recognized by the phospho-Y218 antibody, we assessed phosphorylation by mass spectrometry instead of Western blotting. As shown in **Figure 5C**, PYRP CAR-T cells displayed elevated levels of CD28 Y218 phosphorylation at 10 minutes post-stimulation compared to WT CAR-T cells (P-value = 0.027). To evaluate how this mutation affects cytokine secretion, we cocultured WT and PYRP CAR-T cells with HPAC^WT^ cells for 24 hours and measured cytokine levels in the supernatant. The PYRP mutation led to an approximately two-fold increase in IL-2 production at both the protein (P-value = 0.0009) and RNA levels (P-value = 0.0003) (**Figure 5D-E**), while IFN-γ secretion remained unchanged (**Figure 5F**). We next assessed the anti-tumor efficacy of WT and PYRP CAR-T cells *in vivo* using a pancreatic cancer xenograft model. We injected 1×10^6^ CAR-T via tail vein injection into NSG mice bearing subcutaneous HPAC^WT^ tumors, and monitored tumor growth over time. Both WT and PYRP CAR-T cells initially controlled tumor growth, relative to the untransduced control (UT) group. At later time points, however, differences between the groups became apparent: by day 53, PYRP CAR-T cells maintained significantly stronger tumor control compared with WT CAR-T cells (P-value = 0.0383) (**Figure 5G**).

**Figure 5.**
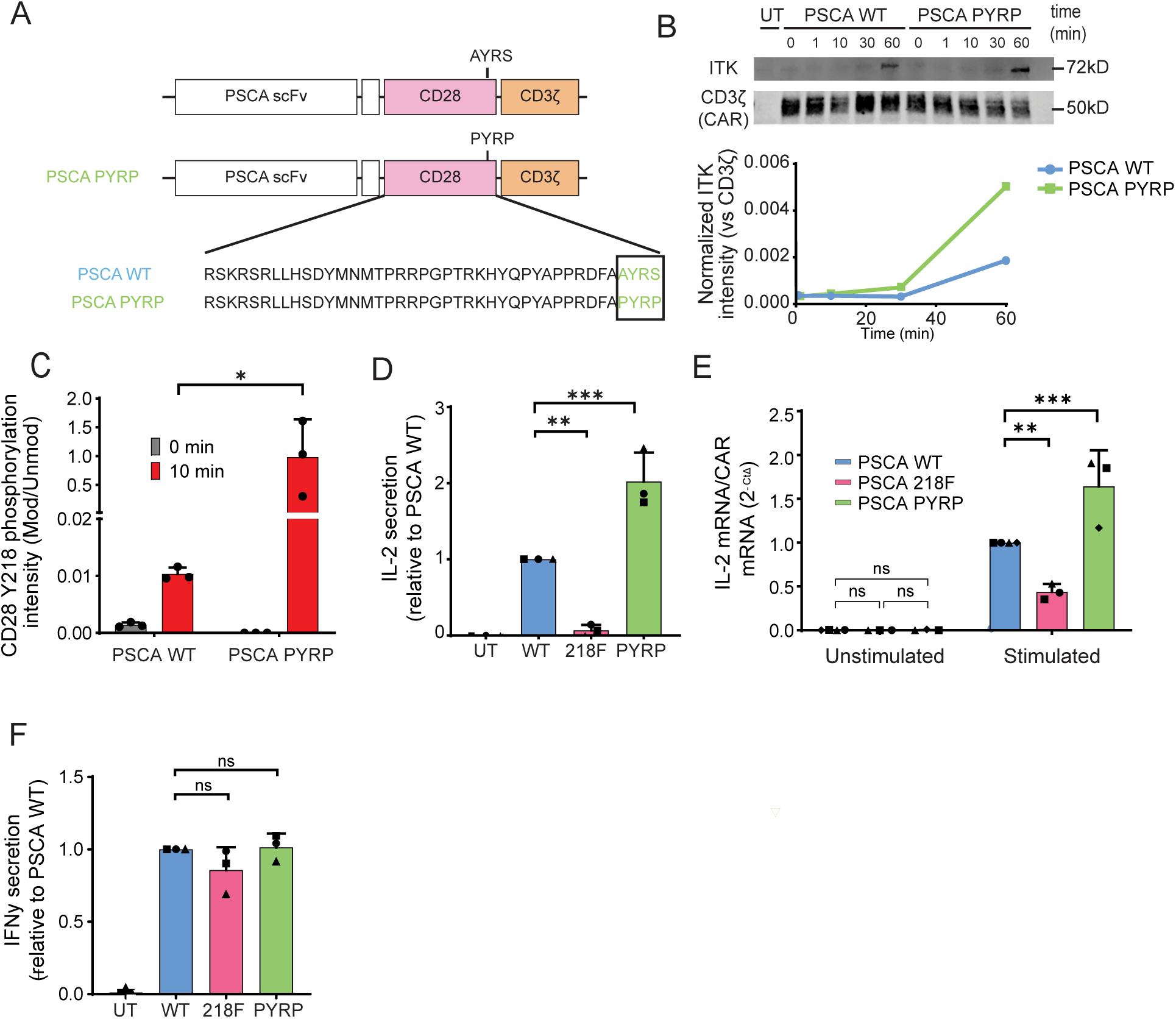
PYRP mutation promotes IL-2 secretion. **A**. Schematic representation of WT and PYRP mutant CAR constructs. In the PYRP mutant, alanine (A) 217 and serine (S) 220 were substituted with prolines (P) to generate a proline-rich domain. **B**. PSCA-targeted WT and PYRP CAR-T cells were cocultured with HPAC^WT^ cells for 0, 1, 10, 30, or 60 min. CAR molecules were immunoprecipitated using protein-L coated beads and co-immunoprecipitated proteins were analyzed by Western blot. Upper panel: Representative Western blot results showing ITK present in CAR immunoprecipitate. CD3ζ staining of the CAR molecule was used as a bait control. Lower panel: Intensity of ITK with respect a total CAR (CD3ζ) was analyzed by densitometry (representative plot). **C.** PSCA WT and PYRP CAR-T cells were stimulated with HPAC^WT^ cells for 10 min and phosphorylation of Y218 was analyzed by mass spectrometry. Bar charts represent the intensity of phosphorylated Y218 normalized to the total CAR signal. Significance was determined by one-way Anova. * = P<0.05. Data is represented as the mean ± standard deviation (SD). **D and F.** PSCA WT and PYRP CAR-T cells were cocultured with HPAC^WT^ cells. Supernatant was collected after 24 hours and cytokine production was measured by ELISA. Significance was determined by one-way ANOVA. * = P<0.05, ** = P<0.01. Each symbol represents an independent experiment from 3 different healthy donors. Data is represented as the mean ± standard deviation (SD). **E.** PSCA WT and PYRP CAR-T cells were cocultured with HPAC^WT^ cells. RNA was extracted after 24 hours and IL-2 mRNA production was evaluated using quantitative Time PCR. CAR mRNA was used as normalization control. Significance was determined by one-way Anova. ** = P<0.01, *** = P<0.001. Each symbol represents an independent experiment from 3 different healthy donors. Data is represented as the mean ± standard deviation (SD). **G.** % of change in tumor volume (compared to baseline) is represented (n=5). Representative example of two independent experiments. Significance was determined by permutation two sample t-test. * = P<0.05.

Altogether, these results highlight the critical role of ITK in CAR-T cell signaling. Our results demonstrate that engineering a synthetic ITK-binding motif into the CAR enhances IL-2 production likely by promoting ITK recruitment and subsequent phosphorylation of CD28 at Y218. In addition, this ITK-binding mutant shows an improved antitumor efficacy *in* vivo compared to WT CAR-T cells suggesting that targeting kinase engagement within CARs can strengthen therapeutic performance..

## DISCUSSION

Chimeric Antigen Receptor T (CAR-T) cell therapy has revolutionized the treatment of hematologic malignancies. However, its efficacy in solid tumors, such as pancreatic cancer, remains limited. Several factors contribute to this challenge, including the immunosuppressive tumor microenvironment, poor infiltration of CAR-T cells into solid tumors, antigen heterogeneity, and T-cell exhaustion ^31^. To overcome these hurdles, multiple strategies have been explored to boost CAR-T cell therapeutic potential. These include the selection of specific T cell subsets ^32,33^, combination therapies with checkpoint inhibitors ^34^, fine-tuning of *in vitro* culture conditions ^35–37^, genetic modifications to regulate exhaustion or activation molecules, such as interleukins or chemokines ^38–40^, and the modification of costimulatory domains ^41^. Common costimulatory domains include CD28 and 4-1BB, each initiating distinct signaling pathways and conferring different metabolic and functional profiles to CAR-T cells. CD28 promotes an effector-like phenotype, enhancing rapid T-cell activation and cytotoxicity, while 4-1BB favors the development of a memory phenotype, contributing to prolonged persistence and sustained activity ^42,43^.

In this paper we have focused on the role of CD28. Despite extensive research throughout the years, the precise signaling pathways mediated by CD28 have been difficult to fully elucidate, revealing complex signaling cascades that often led to conflicting or inconclusive results ^11^. Not surprisingly, this complexity has also become evident during the study of the function of CD28 in the context of CAR-T cell therapy. For instance, some reports have shown that deletion of PYAP binding domain enhances antitumor responses ^44,45^, whereas others have demonstrated that preserving PYAP while mutating YMNM and PRRP improved T-cell function ^46^. Moreover, Guedan et al., showed that introduction of a single amino acid mutation to mimic the ICOS signaling, led to a profound effect in CD28 signaling, persistence and antitumor activity ^47^. Adding to this complexity, we identified a previously underexplored but critical motif within CD28 – tyrosine 218. We demonstrate that mutation in tyrosine 218 in CD28 leads to a severe defect in the antitumor efficacy of CAR-T cells. These findings highlight the importance of the Y218 phosphorylation for CAR-T cell activity. In addition, we hypothesize that the discrepancies observed in the studies of different signaling subdomain might be influenced by the lack of attention given to the last tyrosine of the CD28, which might compensate for the lack of signaling in other mutated subdomains.

The 218F mutant CAR also showed an impaired production of Interleukin-2 (IL-2). This cytokine, crucial for T cell function, is primarily produced by activated CD4^+^ T cells. It plays a key role in promoting T cell proliferation, survival, and differentiation while also regulating immune responses by supporting effector T cell expansion and maintaining immune homeostasis through regulatory T cells (Tregs) ^24^. Our findings demonstrate that reduced IL-2 production is associated with reduced proliferation, a lower overall cell number following chronic stimulation, and lower antitumor efficacy *in vivo*. Together, these results underscore the critical role of Y218 in sustaining CAR-T cell function. It is worth noting that Salter et al. included an 218F mutant in a panel to evaluate LCK binding to the CD28, but reported no functional differences compared to wild-type CAR-T cells. They reported no effect of Y218 mutation on IFN-γ production, which is consistent with our own data; in our case, the functional divergence lies in IL-2 production. Moreover, they reported no change in proliferation, whereas our experiments reveal a measurable effect. We speculate that this effect becomes apparent only in longer experiments, as the differences are subtle and likely detectable only when cumulative, as exemplified by our chronic stimulation assays. These observations suggest that while previous work did not detect a major role for Y218, our more extensive analyses reveal it as a critical regulator of the activity of certain CAR-T cells.

To further investigate the underlying mechanisms behind this observed phenotype, we examined downstream signaling pathways affected by the Y218F mutation and identified PLCγ and SLP76 as key players. Notably, we observed an increase in pSLP76 S376 in the presence of the 218F mutation. SLP76 (SH2 domain–containing leukocyte protein) is a crucial element in TCR-mediated activation of T cells. Phosphorylation of tyrosine residues in this protein, along with LAT, contributes to the formation of a multiprotein complex that facilitates recruitment of various effector molecules and directs signals to downstream pathways. However, SLP76 is also involved in a lesser-known negative feedback loop that dampens the intensity of the TCR signal ^48^. Bartolo *et al*. have demonstrated that HPK1-dependent phosphorylation of Serine 376 in SLP76 also impacts IL-2 production, as its ablation leads to significantly elevated levels of IL-2 secretion. Based on this finding we posit that the higher levels of SLP76 S376 phosphorylation observed in the Y218F mutant may contribute to a lower level of IL-2 secretion. Future studies will be focused on determining the relevance of SLP76 in this pathway.

In addition to impaired IL-2 production, our transcriptomic analysis revealed a notable alteration in the CD4⁺ compartment of 218F CAR-T cells, characterized by upregulation of IL-17A, IL-17F, and related Th17 cytokines, including IL-22 and IL-26. This Th17-skewed transcriptional signature suggests that the 218F mutation promotes CD4⁺ T cell polarization toward a pro-inflammatory Th17-like phenotype. The role of Th17 cells and IL-17 signaling in cancer remains controversial. Several studies have associated IL-17 production with worse prognosis in malignancies such as breast ^49^, colorectal ^50,51^, and lung cancer ^52^, and IL-17 has been implicated in promoting angiogenesis, tumor growth, and metastasis ^53,54^. Consistent with these findings, we observed a correlation between increased IL-17 production and reduced antitumor efficacy of CAR-T cells, although the direct functional contribution of IL-17 to this phenotype remains to be clarified. In contrast to this trend, some studies have reported that adoptive transfer of Th17-polarized T cells can exhibit superior antitumor activity compared to Th1 cells, at least in certain preclinical models such as melanoma. However, these Th17-like cells also exhibited increased IL-2 production relative to their Th1 counterparts—suggesting that IL-2 availability may be critical for the functional capacity of Th17-polarized cells. Based on these observations, we hypothesize that in our model, the combination of impaired IL-2 production—potentially compromising proliferation and persistence—together with a shift toward a Th17-like phenotype and elevated IL-17 production may underlie the reduced antitumor function of 218F CAR-T cells. Although this remains an attractive hypothesis, further studies will be necessary to dissect the individual contributions of IL-17 signaling and IL-2 deficiency to the dysfunctional phenotype observed.

To better understand the molecular basis of CD28 Y218 phosphorylation, we also investigated the upstream regulators of this event. Using both pharmacological inhibition and genetic ablation approaches, we identified ITK as the kinase responsible for this modification. Based on this finding, we hypothesized that promoting ITK binding to the CAR would lead an increased phosphorylation of tyrosine 218 and enhance CAR-T cell function. To test this premise, we generated a novel CAR construct, in which the amino acids surrounding Y218 were mutated to mimic a natural ITK-binding motif (PYRP). As predicted, this mutation led to an increased recruitment of ITK and elevated Y218 phosphorylation compared to the WT CAR. Notably, PYRP CAR-T cells also exhibited enhanced IL-2 secretion.

These results reinforce the idea that Y218 phosphorylation is essential for IL-2 production in CAR-T cells. Prior studies have shown the benefit of combining IL-2 with CAR-T cell therapies. For instance, Sijin Li et al. demonstrated that constitutive IL-2 expression improves anti-tumor activity ^55^. Kagoya et al. designed a CAR construct with antigen-dependent activation of IL-2 related pathways ^56^ and Sockolosky et al. redirected a mutant IL-2 to specifically CAR-T cells ^57^. In line with this model, the PYRP design, with higher IL-2 secretion, also shows improved anti-tumor efficacy *in vivo* compared to the WT CAR.

Overall, our findings provide novel insights into the molecular mechanisms regulating CAR-T cell efficacy in solid tumors. We have shown that the Y218 phosphorylation site within the CD28 costimulatory domain is essential for CAR-T cell anti-tumor effect, and it regulates production of IL-2, a cytokine proven to benefit CAR-T cells performance. In addition, by targeting specific signaling pathways, we demonstrate a potential approach to enhance CAR-T cell performance in challenging tumor environments such as pancreatic cancer.

## MATERIALS AND METHODS

### Generation of Y218 mutant CARs

We used a pMSGV1 backbone for all the plasmids in this study. A PSCA-28t28z CAR (PSCA WT) construct was previously described ^14^. To generate the 218F mutant, the intracellular domains of the PSCA-28t28z CAR was replaced with a gBlock (Integrated DNA Technologies, IA, USA) modified to encode a phenylalanine codon in position 218, through enzymatic digestion (*Not*I/*BamH*I) and ligation. The PYRP mutant construct was generated using the Q5® Site-Directed Mutagenesis Kit (New England Biolabs, E0554S) to substitute Alanine (A) 217 and Serine (S) 220 for Proline (P). Primers used for Site-Directed Mutagenesis: F_PYRP (cgccccAGAGTGAAGTTCAGCAGGAGC) and R_PYRP (atagggTGCGAAGTCGCGTGGTGG). All plasmids were validated through sequencing by Azenta/Genewiz (MA, USA). Please note that amino acid positions are indicated based on wild type CD28 (UniProt P10747) as reference sequence. This nomenclature does not reflect the position of residues within the CAR construct.

### Cell lines

Human Pancreatic Ductal Adeno Carcinoma (HPAC, CRL-2119) cells were obtained from the American Type Culture Collection (ATCC, Manassas, VA). The culture medium consisted of equal volumes of Dulbecco’s modified Eagle’s medium (DMEM) and Ham’s F12 medium containing 2.438 g/L sodium bicarbonate (Gibco, 11320033) supplemented with 2.5 mM GlutaMAX (Gibco, 35050-061), 1X Antibiotic-Antimycotic (Gibco, 15240062), 15 mM HEPES (Corning, 25-060-CI) and 0.5 mM sodium pyruvate (Life Technologies, 11360-070), 0.002 mg/ml insulin (Sigma, I0516), 0.005 mg/ml transferrin (Sigma, T8158), 40 ng/ml hydrocortisone (Sigma Aldrich, 3867), 10 ng/ml mouse epidermal growth factor (SinoBiological, 10605-HNAE), and 5% fetal bovine serum. PSCA deficient HPAC cells (HPAC^PSCA-KO^) were generated by Synthego (CA, USA) and a clonal population was derived by limiting dilution.

Jurkat cells (TIB-152) were obtained from ATCC and cultured in 10% FBS Roswell Park Memorial Institute medium (RPMI, Life Technologies, 61-870-127) supplemented with 2.5 mM GlutaMAX (Gibco, 35050-061), 1X Antibiotic-Antimycotic (Gibco, 15240062). ITK expression was disrupted in Jurkat cells through CRISPR/Cas9 genome editing using a predesigned guide RNA 9gRNA) targeting exon 1 (Hs.Cas9.ITK.1.AA: AAGCGGACTTTAAAGTTCGA Integrated DNA Technologies, IA, USA). Editing was performed using the P3 Primary Cell 4D-Nucleofector X Kit S (Lonza, V4XP-3032) following manufacturer instructions. A clonal cell line of ITK deficient Jurkat cells was generated by limiting dilution.

### Viral production and CAR-T cell transduction

RD114-pseudotyped retroviruses were generated by transient transfection of Human Embryonic Kidney (HEK) 293GP cells as previously described ^13,14^. Healthy donor buffy coats were obtained from OneBlood, Florida Blood Services, FL. Peripheral blood mononuclear cells (PBMCs) were isolated using Ficoll-Paque Plus (GE Healthcare, 17-1440) density gradient centrifugation according to the manufacturer’s instructions and cryopreserved in FBS 10% DMSO. PBMCs were thawed and activated with αCD3 (OKT3) antibody (BioLegend, 317347) at 50ng/ul for 48h. For retroviral transduction, non-tissue culture treated plates were coated with 1.5ml of RetroNectin 20 μg/mL (Takara, T100B) overnight and blocked using 2% BSA PBS for 30 minutes before transduction. In each well, 2×10^6^ T cells were plated in 2ml of complete X-VIVO (Lonza, 04-418Q) alongside with 2ml of thawed retroviral supernatant. Plates were then centrifuged at 2,000 x g at 32°C for 1 hour. The next day 1.7 ml were removed from each well and 2ml of viral supernatant was added. Plates were again centrifuged at 2,000 g at 32°C for 1 hour. After transduction, CAR-T cells were expanded for 5-7 day in X-VIVO (Lonza, 04-418Q) supplemented with 2.5 mM GlutaMAX (Gibco, 35050-061), 1X Antibiotic-Antimycotic (Gibco, 15240062) and 300U/L of IL-2 (MKC RX, NDC 76310-022-01). After expansion CAR expression was assessed by flow cytometry.

### Flow Cytometry

Cells were first washed with PBS and live/dead stained with Fixable dye Near-IR (Thermo, L10119) for 10 minutes at room temperature. For surface staining, cells were incubated with an antibody cocktail diluted in PBS containing 2% BSA for 30 minutes at 4_°C. For intracellular staining (ITK detection), cells were fixed and permeabilized using the Foxp3 Transcription Factor Staining Buffer (eBioscience, 00-5523-00) according to manufacturer instructions. When necessary, CountBright™ Absolute Counting Beads (Thermo Fisher, C36950) were added to enable absolute cell quantification.

The following antibodies were used for immune and cancer cell phenotyping: G4S Linker AF488 (Cell signaling, 50515), CD3 BV711(biolegend, 317328), CD4 PE-Cy7(Biolegend, 300512), CD8 BUV395 (BD, 563795), rabbit ITK (Cell Signaling, 77215), anti-rabbit AF488(Cell Signaling 4412S), PSCA PE (Santa Cruz, sc-80654).

### Cytokine production

To evaluate the ability of CAR-T cells to produce cytokines, we cocultured them with HPAC^WT^ cells at a 1:1 effector-to-target (E:T) ratio for 24 hours in complete RPMI media at 37°C. To independently assess the contribution of CD4⁺ and CD8⁺ cells, these two populations were isolated using human CD4 MicroBeads (Miltenyi, 130-045-101) and human CD8 MicroBeads (Miltenyi, 130-045-201) according to the manufacturer’s instructions. After separation, CD4⁺ and CD8⁺ T cells were mixed at different CD4:CD8 ratios (100:0, 75:25, 50:50, 25:75, and 0:100) and cocultured with HPAC^WT^ cells. Following coculture, the supernatant was collected for cytokine quantification. IL-2 levels were measured using the ELISA MAX™ Deluxe Set Human IL-2 (BioLegend, 431815) following the manufacturer’s protocol. IFNγ levels were measured using ELISA as previously described ^13^. High-binding 96-well plates were coated overnight at 4 °C with anti-human IFNγ capture antibody (Invitrogen, M700A), then blocked with 4% BSA in PBS for 1 hour at room temperature. After washing, samples and standards were added and incubated for 2 hours at room temperature. Bound IFNγ was detected using a biotin-conjugated anti-human IFNγ detection antibody (Invitrogen, M701B), followed by streptavidin-HRP and TMB substrate. The reaction was stopped with 1 M H₂SO₄. IL-2 and IFNγ quantification was performed in technical triplicates and results are representative of at least n=3 independent experiments.

IL-17A secretion was quantified using the Ella™ microfluidic immunoassay platform (ProteinSimple, Bio-Techne) according to the manufacturer’s instructions. Supernatants were collected from coculture supernatants after 24 hours of incubation. Prior to the assay, samples were diluted 1:3 in the Sample Diluent provided with the IL-17A Simple Plex Cartridge (Bio-Techne, SPCKE-PS-009531, V5). Cartridges were loaded with 50 μL of each diluted sample, as well as standard and control reagents, and run on the Ella instrument.

### Repeated Antigen Stimulation

To simulate chronic antigen exposure, CAR-T cells were subjected to repeated *in vitro* stimulation with HPAC^WT^ target cells. CAR-T cells were initially cocultured with HPAC^WT^ cells at a 5:1 E:T ratio in complete RPMI media (without IL-2). After 48 hours, half of the culture was discarded, and the remaining cells were resuspended restimulated with 100.000 newly plated HPAC^WT^ cells. This process was repeated once more at day 4. On day 7, all remaining cells were harvested and quantified by flow cytometry using counting beads (Thermo Fisher, C36950).

### Western blotting and immunoprecipitation

To detect phosphorylation of the CAR signaling domains (CD3 or CD28) or CAR-induced phosphorylation of additional proteins, CAR-T cells were stimulated using HPAC^WT^ cells at a 5:1 ratio at the indicated time points. For non-stimulated samples (0 minutes), CAR-T cells and HPACs were combined after protein extraction. To evaluate the role of ITK in Y218 phosphorylation, ITK inhibitors were incubated with CAR-T cells at 1μM ^15^, for 2 hours, prior stimulation with HPAC^WT^. BMS-509744 and Ibrutinib were purchased from MedChemExpress (HY-11092 and HY-10997 respectively) and GNE-9822 was provided by Genentech under an MTA. To detect Y218 phosphorylation of endogenous CD28, T cells were isolated from healthy donor- derived PBMCs using the Pan T Cell Isolation Kit – human (Miltenyi Biotec, 130-096-535). T cells were then activated using CD3/CD28 T -activator Dynabeads® (Fisher Scientific, 11-131-D) at a 1:1 ratio. For non-stimulated samples (0 minutes), beads were incubated following protein extraction to achieve CD28 binding while avoiding activation.

Cell pellets were collected at indicated time points and protein was extracted using RIPA buffer (Alfa Aesar, J62524-AE) supplemented with protease inhibitor tablets (ThermoFisher, A32953) and phosphatase inhibitor cocktail 2 and 3 (Sigma-Aldrich, P5726 and P0044 respectively). When cell number exceeded 20 million, samples were sonicated using a Bioruptor (Diagenode, 8 pulses of 30 s, with 30 s of rest time). Samples were then centrifuged at 15 500 x g for 45 minutes at 4°C. Protein concentration of the supernatant was determined using the Pierce BCA™ Protein Assay Kit (Thermo-Scientific) following the manufacturer’s instructions.

When specified, CAR was immunoprecipitated using Pierce Protein L Magnetic Beads (Thermo Fisher Scientific, 88849) according to the manufacturer’s instructions. Briefly, magnetic beads were washed with RIPA buffer and then incubated with protein extracts overnight at 4°C with gentle agitation on an orbital mixer. When assessing endogenous CD28, the CD28 protein was enriched by magnetic isolation using CD3/CD28 T-activator Dynabeads®.

Both whole cell extract and immunocomplexes were reduced with 4x Laemmli Sample Buffer (BioRad, 1610747) and heated at 95°C for 10 minutes. Protein was loaded onto a 4–15% Mini-PROTEAN® TGX™ Precast Gel (Bio-Rad, 456-1086) and transferred to a nitrocellulose membrane. Membranes were blocked for 1 hour at room temperature (RT) temp and then incubated for 2 hours at RT or overnight with a primary antibody. Primary antibodies: rabbit anti-CD28 pY218 (Abcam, ab138404), anti-CD28, rabbit ant-ZAP70 pY319 (Cell Signaling, #2717), rabbit anti-ZAP70 (Cell Signaling, #2705), rabbit anti-PLCγ pY783 (Abcam, ab76155), rabbit anti-PLCγ (Abcam, ab76155), rabbit anti-VAV1 pY174 (Abcam, ab76225), rabbit anti-VAV1 (Abcam, ab97574), rabbit anti-LCK (Abcam, ab32149), ITK (Cell Signaling, 77215), mouse anti-CD3ζ (Santa Cruz, sc-166275), mouse anti-B actin (Santa Cruz, sc-47778). Membranes were washed four times in Phosphate Buffer Saline plus 0.1% Tween20 (PBS-T) and incubated with IRDye® 800CW donkey anti-rabbit or IRDye® 680RD donkey-anti-mouse secondary antibodies (LICOR 925-32213 and 925-68072 respectively) for 1 hour at RT. The membranes were washed four times with PBS-T, dried between filter paper and visualized with LI-COR Odyssey imaging system.

### xCELLigence real-time cytotoxicity assay

The cytolytic capacity of mutant CAR-T cells was assed using xCELLigence real-time cytotoxicity assay (RTCA) Multiple Plates instrument (Agilent Technologies, CA, USA). After E-plate calibration 10.000 HPAC^WT^ cells were plated in each well and incubated for 20-24 hours to allow them to adhere. Next, CAR-T cells were added at different ratios and impedance was measured every 15 minutes for 3-4 days. Data analysis was performed with the RTCA software (V 2.0.0.1301.2012, Agilent Technologies, CA, USA). Cell Index was normalized to that of when the T cells were added to the plate and percentage cytolysis was calculated relative to HPAC^WT^ cells grown without effector T cells.

### Proteomics

WT and PYRP CAR-T cells were cocultured with HPAC^WT^ target cells at a 5:1 ratio for 10 minutes. Protein extraction and protein L immunoprecipitation of CAR was performed as described previously. Samples were then separated by SDS-PAGE using a 4–15% TGX gel (Criterion TGX, Bio-Rad, Berkeley, CA) and visualized with Coomassie Brilliant Blue G-250 (Sigma-Aldrich, St. Louis, MO). The bands slightly above 50 kDa were excised for digestion. The gel bands were destained and disulfides were reduced with 2 mM tris(carboxyethyl)phosphine and then cysteines were alkylated with 20 mM iodoacetamide prior to overnight digestion with sequencing grade trypsin (Promega, Madison, WI). The resulting tryptic peptides were extracted with aqueous 50% acetonitrile containing 0.1% trifluoroacetic acid and concentrated by vacuum centrifugation (SC210A, Speedvac, Thermo, San Jose, CA). Peptides were resuspended in 2% acetonitrile with 0.1% formic acid.

Liquid Chromatography-Tandem Mass Spectrometry (LS-MS/MS) discovery proteomics was used to identify the candidate target CAR phosphorylated peptides. An Evosep Ultra High Performance Liquid Chromatography (UHPLC) system coupled to an electrospray orbitrap mass spectrometer (Q-Exactive Plus, Thermo, San Jose, CA) was used for tandem mass spectrometry peptide sequencing. Peptides were separated on an analytical column (EV1106, 15 cm length x 150 µm ID, 1.9 µm particle size) using an extended 88-minute gradient with water + 0.1% formic acid (solvent A) and acetonitrile + 0.1% formic acid (solvent B). The spray voltage was set to 1900 V, the capillary temperature was 275 °C and the S lens RF level was set at 50. MS resolution was set at 70,000 and MS/MS resolution was set at 17,500 with max IT of 50 ms. The top sixteen tandem mass spectra were collected using data-dependent acquisition (DDA) following each survey scan. MS and MS/MS scans were performed with accurate mass measurement using 15 second dynamic exclusion for previously sampled peptide peaks. Sequest ^16^ and Mascot ^17^ searches were performed against the UniProt human database downloaded April 2023 with the WT CAR and PYRP mutant CAR sequences added. Four trypsin missed cleavages were allowed and the precursor mass tolerance was 10 ppm. MS/MS mass tolerance was 0.05 Da. Dynamic modifications included oxidation (M), carbamidomethylation (C), and phosphorylation (S, T and Y).

The peptides identified as candidate targets were monitored in all the samples by parallel reaction monitoring (PRM). LC separation was performed on a Vanquish Neo System (Thermo Scientific) with a C18 reversed-phase analytical column (75 µm ID x 25 cm, 2 µm particle size, 100Å pore size) at 40 °C and 300 nL/minute flow rate. Mobile phase A consisted of 0.1% formic acid in water. Mobile phase B consisted of 0.1% formic acid in 100% acetonitrile. The 105-minute gradient was programmed as follows; solvent B from 1% to 2% in 5 minutes, then solvent B from 2% to 30% B in 80 minutes, followed by solvent B from 30% to 38.5% B in 10 minutes, then solvent B from 38.5% to 90% B in 5 minutes and held at 90% for 5 minutes. Eluting peptides were ionized by electrospray ionization and then analyzed by an Orbitrap Ascend Tribrid mass spectrometer (tune version 4.0.4091, Thermo Scientific) with high field asymmetric waveform ion mobility spectrometry (FAIMS) using PRM. Spray voltage was set to 2 kV. Ion transfer tube temperature was set to 305 °C. RF lens was set to 50. Two FAIMS compensation voltage values (-45 and -65) were used with 2 second cycle time each for each acquisition. PRM analysis was unscheduled. MS data was not acquired in these experiments. MS/MS resolution was 60,000. and MS/MS resolution was 60,000. PRM scans were set up to target both the phosphorylated and unmodified candidate peptides (WT CAR peptides, DFAAYR and DFAAYR, and PYRP mutant CAR peptides, DFAPYRPR and DFAPYRPR).

Skyline was used for peak integration of PRM data ^18^. The PRM Skyline document initially scanned for all b and y fragment ions from ion 2 (i.e. b2 and y2) to the next to the last ion in both series (i.e. b(n-2) and y(n-2), where n is the number of amino acids in the peptide). The document was then refined to include only the top 4-5 ions for each peptide, favoring larger m/z fragment ions to reduce potential interference. Each peak was manually assessed, and the peptide peak area was exported. The peak area of the modified WT CAR phosphopeptide and PYRP mutant CAR phosphopeptide from each sample was normalized to its respective unmodified peptide.

### Animal experiments

All animal studies were performed in compliance with institutional regulations on animal use. Procedures were approved by the University of South Florida Institutional Animal Care and Use Committee (IACUC, protocol R IS00002385). Experiments were performed in the Comparative Medicine Facility. Six to eight week-old male NSG 5557 mice were purchased from the Jackson Laboratory (Bar Harbor, ME). Xenograft tumors were generated by subcutaneous injection of 2×10^6^ HPAC^WT^ cells in PBS. Two weeks after tumor cell injection, mice were treated with intravenous injection of 5×10^6^ T cells. Tumor dimensions were assessed twice a week using calipers. Tumor volume was calculated using the formula 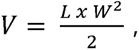, where L is the length and W is the width.

### RNA-seq and bioinformatics analysis

WT and 218F CAR-T cells were stimulated using HPAC^WT^ cells at 5:1 ratio for 24h. After incubation CD4^+^ and CD8^+^ cells were labeled using fluorescent antibodies and sorted using MACSQuant® Tyto®. Because of the nature of this instrument, sequential sorting was necessary. Briefly, CD4^+^ T cells were first sorted, and the negative fraction - containing CD8^+^ cells and tumor cells - was reloaded in the cartridge to sort for the CD8^+^ cells only. To prevent cell damage, sorted cells were immediately washed and resuspended in RLT buffer. RNA isolation was performed using Qiagen RNeasyMini Kit (Qiagen, cat. #74104) following manufacturer’s instructions.

RNA was quantitated with the Qubit Fluorometer (ThermoFisher Scientific, Waltham, MA) and screened for quality on the Agilent TapeStation 4200 (Agilent Technologies, Santa Clara, CA). The samples were then processed for bulk RNA-sequencing using the NuGEN Universal Plus Human RNA-Seq Library Preparation Kit with NuQuant (Tecan Genomics, Redwood City, CA). Briefly, 100 ng of RNA was used to generate cDNA and a strand-specific library following the manufacturer’s protocol. Quality control steps were performed, including TapeStation size assessment and quantification using the Kapa Library Quantification Kit (Roche, Wilmington, MA). The final libraries were normalized, denatured, and sequenced on the Illumina NovaSeq 6000 sequencer with the SP-200 cycle reagent kit in order to generate approximately 45M million 100-base read pairs per sample (Illumina, Inc., San Diego, CA).

FASTQ files were aligned to the Homo sapiens (human) genome assembly GRCh38 (hg38, Genome Reference Consortium, GCA_000001405.15 GCF_000001405.26) reference genome using STAR aligner ^19^ with the default setting. Gene quantification was performed using HTSeq ^20^. Custom scrips were developed for analysis. DESeq2 ^21^ was used to determine differentially expressed genes between CD4^+^ and CD8^+^ WT and 218F CAR-T cells. The enriched pathways were analyzed through the use of QIAGEN IPA (QIAGEN Inc., https://digitalinsights.qiagen.com/IPA) ^22^.

### Statistical Analyses

Statistical analyses were performed using GraphPad Prism version 10.2.3 and R version 4.2.3. For *in vitro* studies, each experiment was repeated independently, with at least three replicates as indicated in the figure legends. Unless otherwise indicated, data is presented as mean ± SD (standard deviation). Comparisons between two groups were assessed using paired or unpaired Student’s t-test. In experiments where donor variability was high, the test group was normalized to the control group (PSCA WT), and statistical significance was assessed by t-test with Welch’s correction to account for unequal variances. For comparisons involving more than two groups, one-way ANOVA followed by multiple comparisons test was applied. For *in vivo* studies, each experiment was repeated independently with two replicates. Tumor growth is expressed as the percentage change in tumor volume from the baseline measurement. Statistical differences between test groups at the final time point were evaluated using a permutation two-sample t-test. For all analysis, a p-value < 0.05 was considered statistically significant

## ACKNOWLEDGEMENTS

This work has been supported, in part, by the Proteomics, Molecular Genomics, and Flow Cytometry Core Facilities at the H. Lee Moffitt Cancer Center and Research Institute, an NCI-designated Comprehensive Cancer Center (P30-CA076292). Additional support was provided by the NCI (R21CA280233) and by the ‘’la Caixa’’ Fellowship Program awarded to E.M-P. This work was also partially supported by a generous donation from the Steinman Family Foundation. We thank Genentech for generously providing the GNE-9822 inhibitor. We are also grateful to Darwin Chang for his assistance with the analysis of the genomics data.

## AUTHOR CONTRIBUTION

E.M-P, M.C.R, M.G.F and D.A-D. designed experiments, E.M-P, M.C.R and M-R. S. performed experiments, L.D. and J.K. performed phospho-proteomics experiment and data analysis. E.M-P, D.A-D. and Y.K. analyzed data and performed statistical analysis. E. M-P. and D.A-D. wrote the manuscript.

## DECLARATION OF INTEREST

D.A-D. and E.M-P. are inventors in a provisional patent application filed by the Moffitt Cancer Center, related to the technology described in this manuscript.

## FUNDING

This work has been supported, in part, by the Proteomics, Molecular Genomics, and Flow Cytometry Core Facilities at the H. Lee Moffitt Cancer Center and Research Institute, an NCI-designated Comprehensive Cancer Center (P30-CA076292). Additional support was provided by the NCI (R21CA280233) and by the ‘’la Caixa’’ Fellowship Program awarded to E.M-P. This work was also partially supported by a generous donation from the Steinman Family Foundation. Sponsors were not involved in the study design, collection, analysis or interpretation of the data or decision to submit the paper for publication.

During the preparation of this work the authors used OpenAI (ChatGPT) in order to improve grammar and enhance the readability of the manuscript. The tool was used strictly for linguistic editing and not for the generation of original ideas or data. After using this tool, the authors reviewed and edited the content as needed and take full responsibility for the content of the publication.

## Supplemetary Figures

**Supplementary Figure 1.**
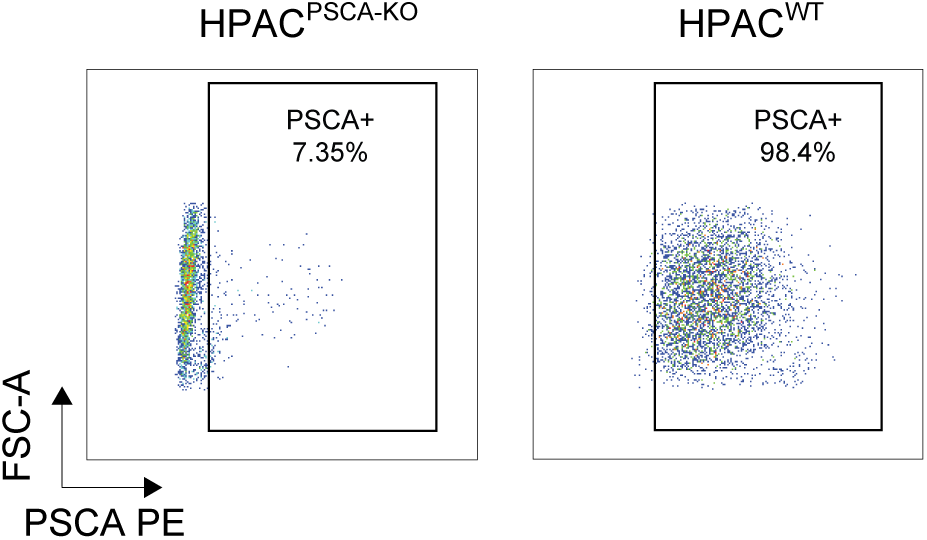
Expression of PSCA in HPAC^WT^ and HPAC^PSCA-KO^ cells, measured by flow cytometry.

**Supplementary Figure 2.**
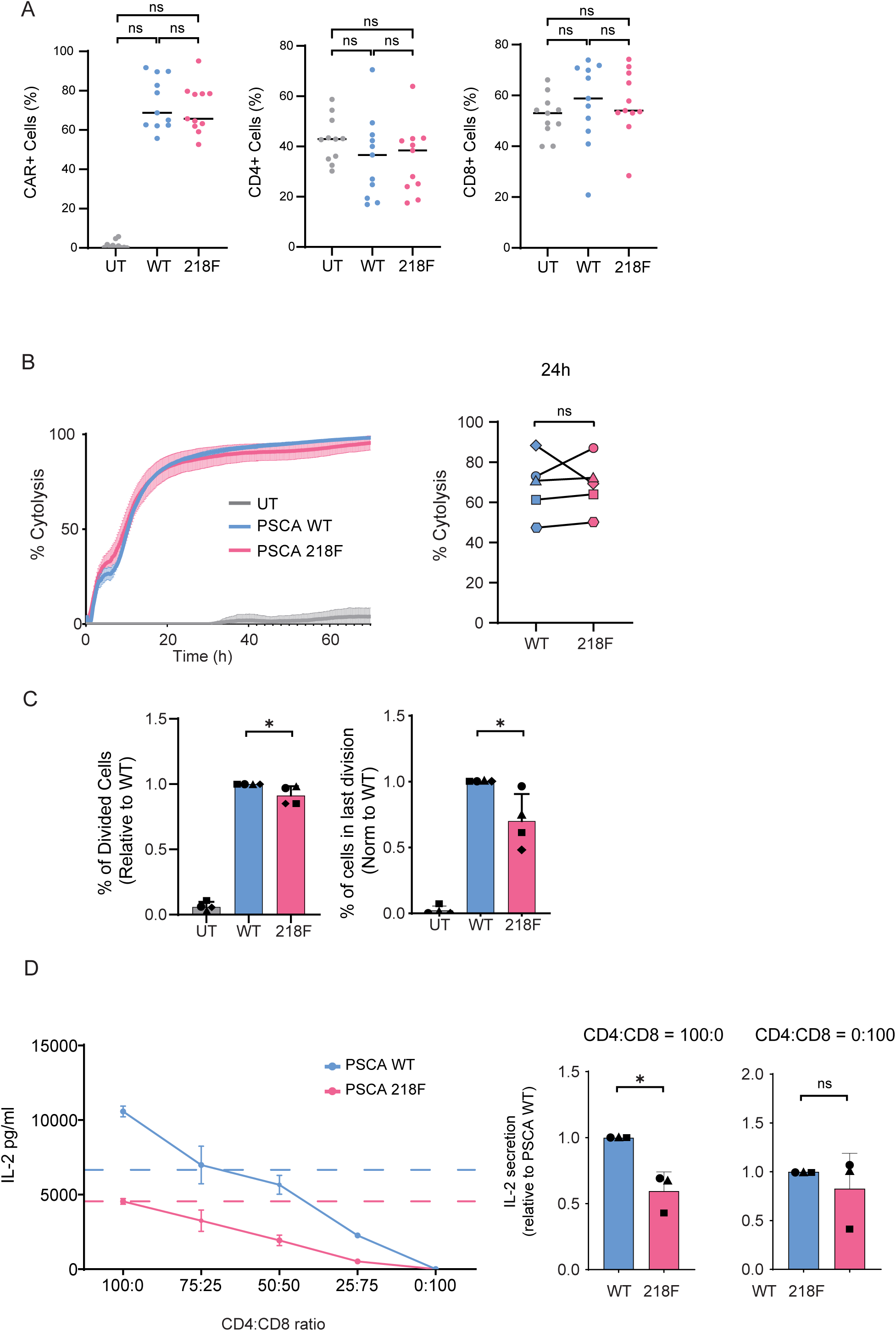
**A.** Expression of CAR, CD4, and CD8 in WT and 218F CAR-T cells 7-10 days after transduction. Each point represents an individual healthy donor (n = 11). Statistical significance was determined by one-way ANOVA. **B.** The lysis of HPAC^WT^ cells when treated with WT or 218F CAR-T cells was evaluated using a real-time cytotoxicity assay (xCELLigence). Left Panel: Representative graph showing % of cytolysis over a 72 hour period after the addition of the effector cells. Right Panel: % of cytolysis of WT and 218F CAR-T cells at 12 hours after the addition of the effector cells. Significance was determined by paired t-test. Each symbol represents an independent experiment using one of five different healthy donors. **C.** Proliferation capacity of WT vs 218F CAR-T cells, analyzed by CellTrace Violet dilution. WT and 218F CAR-T cells were co-cultured at a 2:1 E:T ratio and incubated at 37°C for 4 days. Percentage of Divided cells (left) and percentage of cells in the last division (right) of 218F CAR-T cells is shown normalized to WT CAR-T cells. Significance was determined by t-test with Welch’s correction. * = P≤0.05. Each symbol represents an independent experiment from 4 different healthy donors. Data is represented as the mean ± standard deviation (SD). **D.** CD4 and CD8 CAR-T cells were sorted using magnetic MicroBeads. WT and 218F CAR-T cells were cocultured with HPAC^WT^ cells at different CD4:CD8 ratios. Supernatant was collected after 24 hours and cytokine production was measured by ELISA. Left Panel. Representative graph showing the IL-2 production at different CD4:CD8 ratio. Dotted lines indicate IL-2 production by unsorted CAR-T cells. Center and left panel show production of IL-2 production of 218F CAR-T cells with respect to WT CAR-T cells. Significance was determined by t-test with Welch’s correction. ** = P<0.01. Each symbol represents an independent experiment from 3 different healthy donors.

**Supplementary Figure 3.**
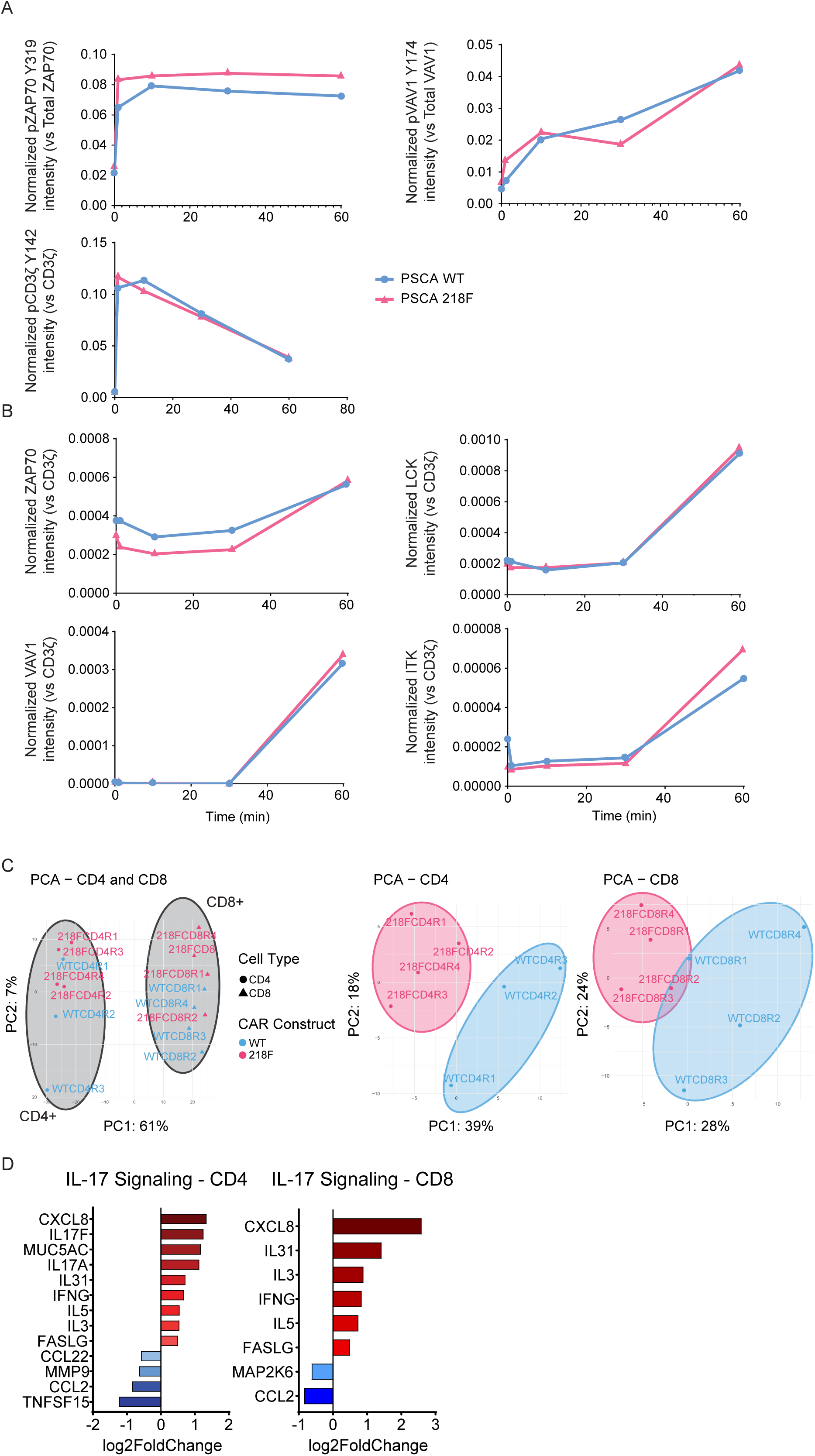
Transcriptomic profile of WT vs 218F CAR-T cells. CAR-T cells were cocultured with HPAC^WT^ cells for 24 hours and subsequently sorted into CD4⁺ and CD8⁺ subsets using the MACSQuant® Tyto® system prior to RNA extraction. **A.** Principal component analysis (PCA) of WT and 218F CAR-T cells in all samples (left), CD4⁺ cells (center), and CD8⁺ cells (right). **B.** Gene expression of genes in the IL-17 signaling pathway. Enriched pathways between WT and 218F CAR-T cells were identified using IPA® software. Bar plots show log₂ fold change in gene expression in 218F CAR-T cells relative to WT CAR-T cells. CD4⁺ cells are shown on the left and CD8⁺ cells on the right.

**Supplementary Figure 4.**
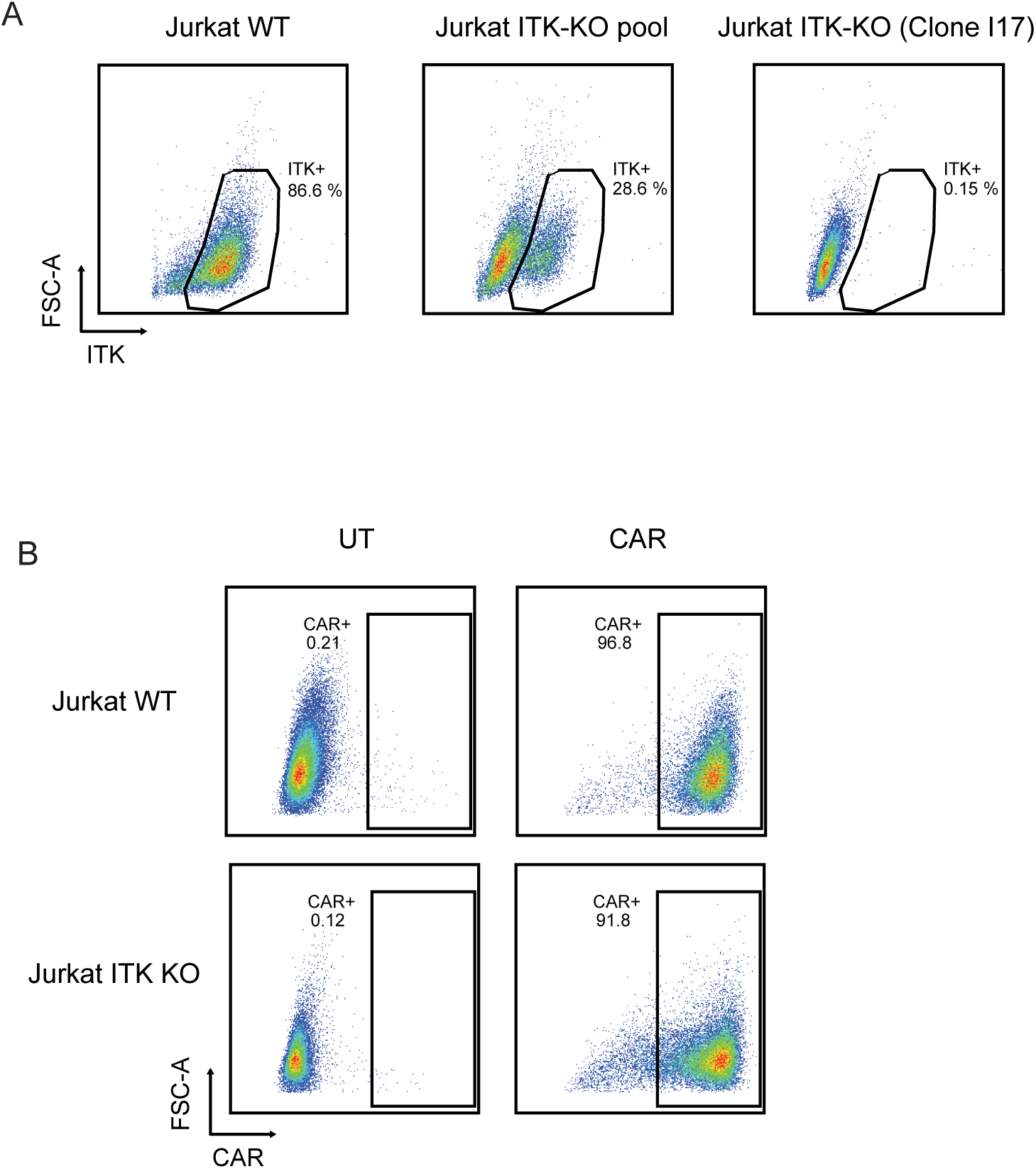
**A.** Expression of ITK in Jurkat cells. ITK-KO Jurkat cells were generated using the CRISPR/Cas9 technology to target exon 1 of ITK. A clonal population was derived by limiting dilution. **B.** CAR transduction efficiency in Jurkat WT and Jurkat ITK-KO 7 days after transduction. Representative plot of one transduction.

